# Alternative Conformations and Motions Adopted by 30S Ribosomal Subunits Visualized by Cryo-Electron Microscopy

**DOI:** 10.1101/2020.03.21.001677

**Authors:** Dushyant Jahagirdar, Vikash Jha, Kaustuv Basu, Josue Gomez-Blanco, Javier Vargas, Joaquin Ortega

## Abstract

It is only after recent advances in cryo-electron microscopy that is now possible to describe at high resolution structures of large macromolecules that do not crystalize. Purified 30S subunits interconvert between the “active” and “inactive” conformations. The active conformation was described by crystallography in the early 2000s, but the structure of the inactive form at high resolution remains unsolved. Here we used cryo-electron microscopy to obtain the structure of the inactive conformation of the 30S subunit to 3.6Å resolution and study its motions. In the inactive conformation, three nucleotides at the 3’ end of the 16S rRNA cause the region of helix 44 forming the decoding center to adopt an unlatched conformation and the 3’ end of the 16S rRNA positions similarly to the mRNA during translation. Incubation of inactive 30S subunits at 42 °C reverts these structural changes. The position adopted by helix 44 dictates the most prominent motions of the 30S subunit. We found that extended exposures to low magnesium concentrations induces unfolding of large rRNA structural domains. The air-water interface to which ribosome subuints are exposed during sample preparation also peel off some ribosomal proteins. Overall this study provides new insights about the conformational space explored by the 30S ribosomal subunit when the ribosomal particles are free in solution.

## INTRODUCTION

Ribosomes in bacteria undergo constant conformational changes that are essential for the translation process. These include from small-scale base flipping events at the decoding center to much large-scale motions of ribosomal subunit domains induced by mRNA, tRNA and translation factor binding (Frank, 2017). Similarly, the process of ribosome assembly involves constant conformational changes, as the rRNA folds and ribosomal proteins are incorporated into the assembling particle (Mulder et al., 2010; Razi et al., 2017a; Sashital et al., 2014). Until recently, high-resolution structural information about these conformational states was solely contributed through X-ray crystallography. Consequently, only those states that could be stabilized in a crystal lattice were accessible providing only a reduced breath of the conformational heterogeneity existing in these processes and potentially masking important details about local and global conformational dynamics.

Today, structural biology is in the midst of a “resolution revolution” (Kuhlbrandt, 2014). Due to continuous advances, cryo-electron microscopy (cryo-EM) can now routinely contribute high resolution models of large macromolecular machines with dynamic composition and conformations that have remained impervious to crystallization (Cheng, 2015; Cheng et al., 2017; Nogales and Scheres, 2015). In the context of the ribosome, these cryo-EM models are illuminating new relevant transition steps in the protein translation (Hussain et al., 2016) and ribosome assembly processes (Ni et al., 2016; Nikolay et al., 2018; Razi et al., 2019; Seffouh et al., 2019).

Particularly, in the study of the ribosome assembly process structural biologist have widely used genetic approaches to trigger accumulation of assembly intermediates (Razi et al., 2017a; Stokes and Brown, 2015). The essence of this approach consists of creating single deletion or depletion strains for one of the assembly factors to disable or slow down the ribosome biogenesis process (Daigle and Brown, 2004). Invariably, these methods produce a heterogeneous mixture of immature ribosomal particles from which is not possible to produce crystals. However, structural characterization of these assembly intermediates using cryo-EM combined with image classification approaches has shown to be a powerful approach to identify the role of protein factors in assisting specific steps in the ribosome assembly process. For example, immature 30S subunits purified from Δ*yjeQ* (Jomaa et al., 2011) and Δ*rimM* (Guo et al., 2013; Leong et al., 2013), two non-essential assembly factors, exhibited structural deficiencies in the decoding region when compared to the mature 30S subunit. Similarly, depletion of Era, an essential assembly factor causes accumulation of 30S subunit presenting unfolding in important functional motifs on the platform region and decoding center (Razi et al., 2019). The conclusion from these studies was that YjeQ, RimM and Era participate in the assembly steps maturing the functional core of the 30S subunit.

Inferring the role of assembly factors from the structural deficiencies observed in the assembling particles accumulating in the null or depleted cells requires the existence of a common standard or reference structure that is used in the comparative analysis. Previous publications have typically used the structure of the mature 30S subunit obtained by crystallographic approaches as the reference structure (Wimberly et al., 2000). However, the mature 30S subunit not constrained on a crystal lattice is not existing only in one conformation. Five decades ago, Elson and colleagues (Zamir et al., 1969, 1971) already reported that purified 30S subunits readily interconvert between “active” and “inactive” conformations. Later Noller’s group determined using chemical probing that transition between both states involves structural changes in the neck and decoding center regions of the 16S rRNA (Moazed et al., 1986). This conformational variability of the mature 30S subunit is not restricted to the bacterial ribosome. The eukaryotic 40S subunit also seems to sample multiple conformations (Swiatkowska et al., 2012). More recently, it was found using RNA SHAPE (selective 2-hydroxyl acylation analyzed by primer extension) that in exponentially growing *Escherichia coli* cells, 16S rRNA largely adopts the inactive conformation in free mature 30S subunits and the active conformation in translating 70S ribosomes (McGinnis et al., 2015; McGinnis and Weeks, 2014). The reactivity patterns on the 30S subunit associated with the 50S subunits are fully consistent with the RNA secondary structure exhibited by the 30S subunit in the crystal structure suggesting that both approaches are describing the same structure. However, the high-resolution structure of the inactive conformation observed in the free 30S subunits has never been obtained by X-ray crystallography or cryo-EM.

To visualize the 3D structure of the inactive conformation of the 30S subunit, and potentially other alternative conformations mature 30S subunits adopt outside the constrains of a crystal lattice, we exposed purified 30S subunits to buffer conditions that recall those in ribosome purification approaches (Daigle and Brown, 2004; Jomaa et al., 2011). Cryo-EM revealed that the mature 30S subunits in solution adopt a variety of conformations. Magnesium concentration in the purification buffers had a large effect. Exposure to low magnesium concentrations switched the decoding center of the 30S subunit to a drastically different conformation from that observed in the crystal structure. Incubation of the purified 30S subunits at 42 °C induced the decoding center to switch back to the canonical conformation. Extended exposure to low magnesium concentration increased the heterogeneity and led to the appearance of particle populations with complete unfolding of the head domain and partial unfolding of the platform domain. Our experiments also showed how the water-air interface to which the 30S subunits are exposed during the vitrification process in cryo-EM can also induce structural variability by causing the loss of r-proteins. Studying the motion of these ribosomal particles revealed that the conformation adopted by helix 44 dictates the main motions exhibited by the different 30S subunit populations.

## RESULTS

### High-resolution cryo-EM structure of the 30S subunit inactive conformation

In actively growing *E. coli* cells only between 2-5% of the existing 30S subunits are in an immature state and remain dissociated without forming 70S ribosomes (Leong et al., 2013; Thurlow et al., 2016). Consequently, 30S subunit purification protocols typically include a step that exposes ribosomes temporarily to buffers containing 1-2 mM concentration of magnesium ions (low magnesium). This condition induces a dissociation of the ribosome into its two integrating subunits, the 30S and the 50S subunits. This is mainly triggered by a structural change in the 30S subunit termed ‘inactivation’, as this conformational change interferes with tRNA binding in its P site (Moazed et al., 1986; Zamir et al., 1969).

To obtain the cryo-EM structure of the inactive conformation of the 30S subunit at high resolution, we purified 30S subunits from actively growing *E. coli* cells using a protocol that exposed the ribosomes to 1.1 mM magnesium acetate. However, at the end of the purification, the buffer in the fractions containing purified 30S subunits was exchanged and the concentration of magnesium acetate raised to 10 mM. The obtained sample (30S-Inactivated-high-Mg^2+^) was then imaged by cryo-EM.

Using image classification approaches, we found that the purified 30S-Inactivated-high-Mg^2+^ subunits existed in two distinct conformations that we called class A and B (Fig. 1A). The distribution of particles between these two classes was 79% and 24%, respectively. Using these particle populations, we calculated a 3.6 Å resolution cryo-EM map for class A and a 4.4 Å resolution map for class B (Supplementary Fig. S1). The structures of the body (5’ domain), platform (central domain) and head domains (3’ major domain) in both structures were identical to the canonical structure of the 30S subunit obtained by X-ray crystallography (Wimberly et al., 2000) (Fig. 1A). However, the decoding region located at the convergency point of all these three domains presented important differences in both maps (Fig. 1B). In the canonical structure, helix 44 runs from the bottom of the body to the lower part of the head. The upper region of this helix near the platform domain is involved in the decoding process and is also a critical element in creating the subunit interface with the 50S subunit. In the map for class A, the upper domain of helix 44 adopts an alternative conformation and is not latched to the decoding center as described by the crystal structure. Instead, this entire section of the helix protrudes from the surface of the 30S subunit and distorts the interface with the 50S subunit. In class B, the entire helix 44 seemed to adopt a flexible conformation. The lower part of this helix showed a well-defined density, but the middle and upper region exhibited a highly fragmented density indicating this region is highly flexible.

**Figure 1.**
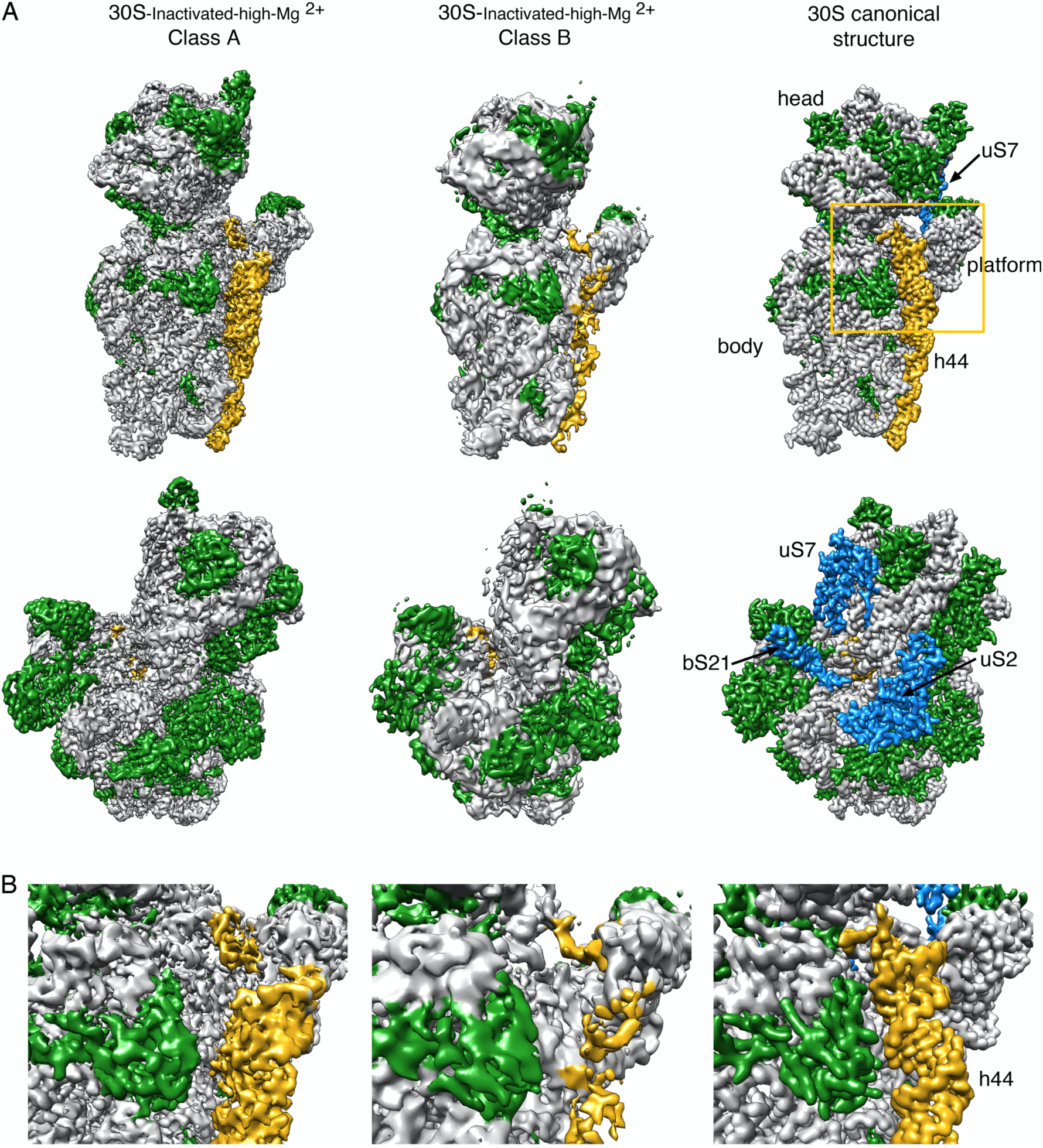
Cryo-EM structure of the 30S-Inactivated-high-Mg^2+^ particle. (A) Front (top row) and back view (bottom row) of the cryo-EM maps obtained for the two subpopulations found for the 30S-Inactivated-high-Mg^2+^ particle. The maps are shown side-by-side with the structure of the 30S subunit obtained by X-ray crystallography (30S canonical structure). This structure was obtained by generating a density map from PDB file 4V4Q and subsequently low pass filtering this structure to 4 Å. The rRNA is displayed in light grey, the r-proteins in green and helix 44 in golden rod orange. The r-proteins uS2, uS7 and bS21 for which a representative density does not appear in the cryo-EM maps of the 30S-Inactivated-high-Mg^2+^ particle are shown in blue in the map of the 30S subunit obtained by X-ray crystallography. These proteins and other landmarks of the 30S subunit are labeled. (B) Zoomed in view of the decoding region of the cryo-EM maps obtained for the two subpopulations found for the 30S-Inactivated-high-Mg^2+^ particle and the structure of the 30S subunit obtained by X-ray crystallography. The area visualized in this panel is indicated as a frame in panel (A).

To quantitatively measure the structural differences in the 30S-Inactivated-high-Mg^2+^ class A with respect to the conventional structure, we produced a molecular model from this map and subsequently calculated a temperature map to measure the deviation of the structure adopted by the 16S rRNA in these ribosomal particles (Fig. 2A). The body and platform regions were highly similar and the rRNA in these structural motifs mostly overlapped with the canonical structure. In contrast, the position of the head and upper domain of helix 44 and contacting region in helix 28 diverged significantly. The head domain was tilted backwards by 16°, opening up the decoding region (Fig. 2B). The upper domain of helix 44 and bottom part of helix 28 adopted a drastically different conformation that was stabilized by a different arrangement in the base paring of the rRNA (Fig. 2C). The transition between both conformations involves nucleotides 1532-1534 at the 3’ end of the 16S rRNA. These three nucleotides are not forming any base pairing in the conventional structure, however in class A they approach helix 28 and unfold its bottom part (region formed by nucleotides 1391-1396 and 921-925) and base pair with nucleotides 921-923 in that region. This transition also causes the partial unfolding of the top of helix 44 formed in the canonical structure by nucleotides 1397-1407 and 1494-1503, as well as the positioning of the 3’ end of the 16S rRNA (distal to nucleotide 1534) in a conformation similar to that adopted by mRNA during translation. In this conformational transition helix 45 slightly shift in position but remains folded (Supplementary Movie 1).

**Figure 2.**
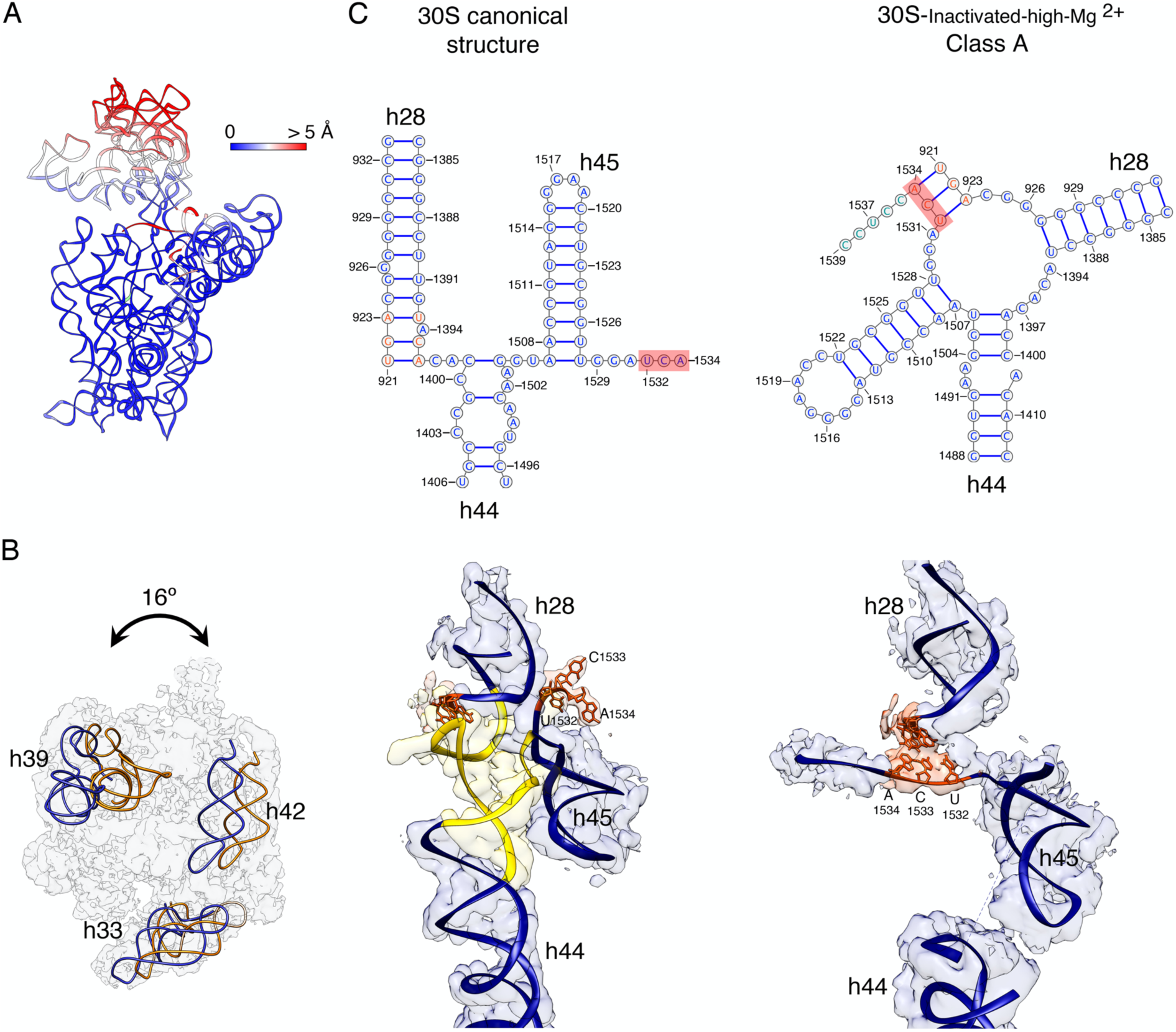
Molecular model of the 30S-Inactivated-high-Mg^2+^ particle. (A) Temperature map of the 30S-Inactivated-high-Mg^2+^ class A. The rRNA is colored according the r.m.s.d. deviation (Å) with respect to the structure of the 30S subunit obtained by X-ray crystallography (PDB ID 4V4Q). (B) Top view of the head of the cryo-EM map of the 30S-Inactivated-high-Mg^2+^ class A. The positions of helices 33, 39 and 42 in this structure (navy blue) and in the crystal structure of the 30S subunit (orange) are shown to illustrate the backwards tilting of the head by 16° in the structure of the 30S-Inactivated-high-Mg^2+^ class A. (C) Secondary (top panels) and tertiary (bottom panels) structures of helices 28, 44 and 45 of the 16S rRNA in the 30S subunit structure obtained by X-ray crystallography and in the molecular model derived from the cryo-EM map of the 30S-Inactivated-high-Mg^2+^ class A. The nucleotides 1532-1534 at the 3’ end of the 16S rRNA involved in the conformational transition between both structures are highlighted in red. The regions of helices 44 and 28 that become unfolded during the conformational transition is colored in yellow in the canonical 3D structure of the 30S subunit.

Taken together, these structures indicate that 30S subunits purified through approaches that expose them to low magnesium concentration switch from the conventional structure shown by X-ray crystallography to an inactive conformation that exhibit drastic structural differences in the decoding region. The inactive conformation is stabilized by an alternative base pairing of the nucleotides in the 3’ end of the 16S rRNA molecule.

### Exposure of the 30S subunits to the air-water interface in the cryo-EM grid causes the loss of r-protein uS2

Previously reported cryo-EM structures of mature 30S subunits (Datta et al., 2007; Razi et al., 2019) showed either fragmented or absent densities for r-proteins uS7 and bS21 suggesting that they are intrinsically flexible when the particle is not constrained in a crystal lattice. Consistently, the densities corresponding to these two r-proteins in the cryo-EM maps obtained for the 30S-Inactivated-high-Mg^2+^ subunit class A and B were also not present (Fig. 1A). Surprisingly, we also found the density representing uS2 to be completely absent from our cryo-EM maps. In the 30S subunit, uS2 binds in the solvent face (convex face) of the subunit stably anchoring its two domains to the 16S rRNA and typically is fully visible in previously obtained X-ray (Schureck et al., 2016; Wimberly et al., 2000) and cryo-EM (Lopez-Alonso et al., 2017; Razi et al., 2017b) structures.

In cryo-EM, to preserve the specimen in their hydrated state, one spread the sample in a thin layer of buffer solution supported in the cryo-EM grid right before plunging the grid into liquid ethane to freeze the liquid layer into vitreous ice. In the vitrification device used in our experiments, the time that elapses between the blotting of the grid to form the thin layer and the vitrification is typically 1 second. In this time, ribosomal particles collide with the air-water interface between 100-1,000 times (Noble et al., 2018). These interactions have the potential to cause damages in the specimen.

To investigate whether the repeated interaction of the ribosomal with the air-water interface was causing the loss of uS2, we repeated the imaging of the 30S-Inactivated-high-Mg^2+^ particles by adding an additional layer of continuous carbon to the grids (Fig. 3A). We hypothesized that by adsorbing the particles to the support film, their exposure to the air-water interface would be reduced. We collected a cryo-EM dataset from these grids and particle images were subjected to a similar image classification workflow. We found that particles were present mainly as one class with helix 44 in an identical conformation to the structure of the 30S-Inactivated-high-Mg^2+^ class A particles. The cryo-EM map obtained from these particles (30S-Inactivated-Carbon-high-Mg^2+^) refined to a resolution of 3.8 Å (Fig. 3B and Supplementary Fig. S1). More importantly, the density representing uS2 was clearly visible in the cryo-EM map.

**Figure 3.**
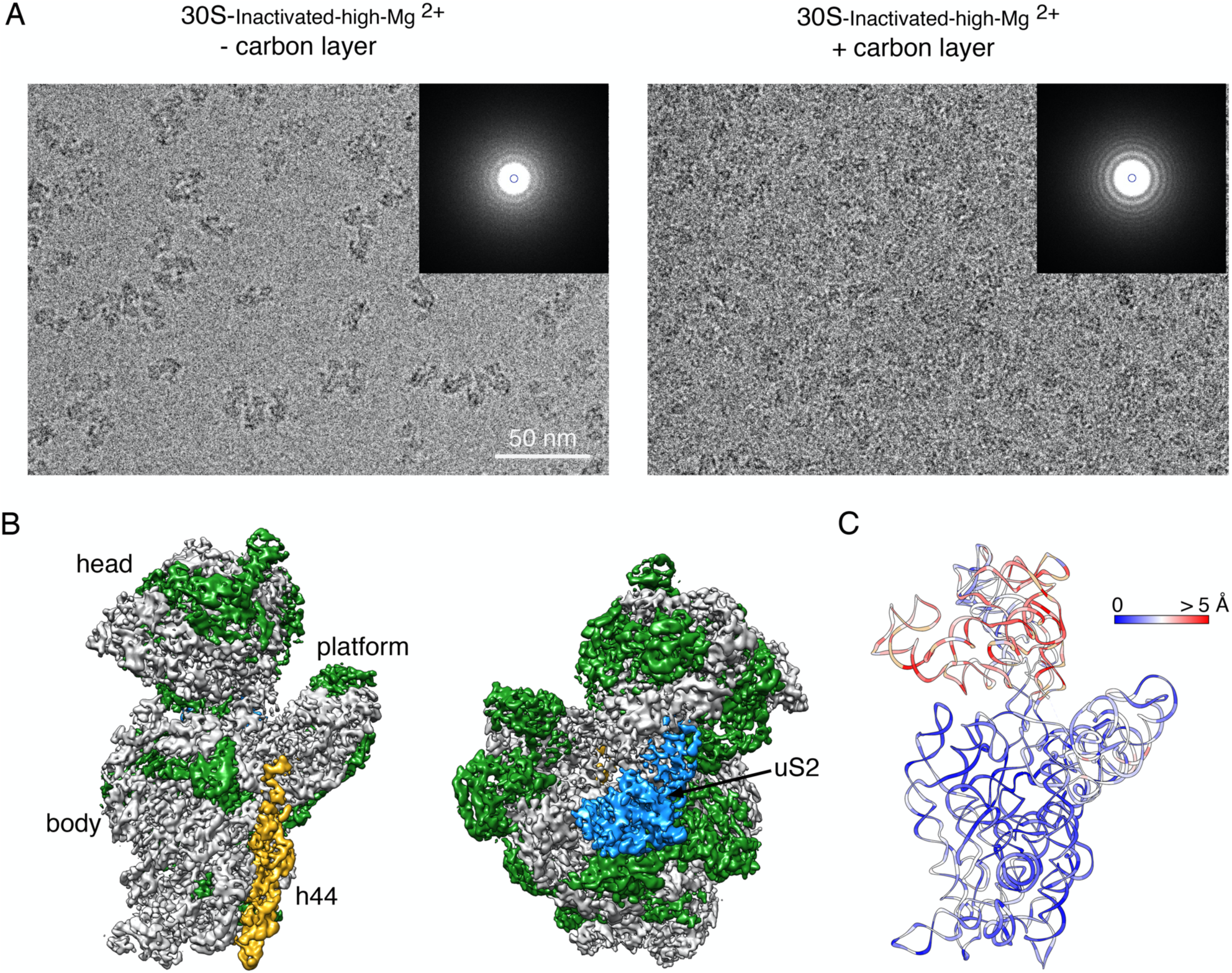
Cryo-EM structure of the 30S-Inactivated-high-Mg^2+^ particle in grids containing a continuous layer of carbon. (A) Two representative electron micrographs containing 30S-Inactivated-high-Mg^2+^ particles. The micrographs were obtained from EM grids without (left panel) and with (right panel) an additional layer of a continuous carbon. The inset shows the power spectra from each micrograph. The presence of a continuous layer carbon makes the Thon rings in the power spectra more prominent and the background of the micrograph more prominent. (B) Front and back view of the cryo-EM map obtained for the 30S-Inactivated-high-Mg^2+^ particle from grids containing a continuous carbon layer. The rRNA is shown in light grey and the r-proteins are shown in green except uS2 that is colored in blue. (C) Temperature map of the 30S-Inactivated-high-Mg^2+^ molecular model obtained from grids containing a continuous carbon layer. The rRNA is colored according the r.m.s.d. deviation (Å) with respect to the structure 30S-Inactivated-high-Mg^2+^ class A obtained from grids without a continuous carbon layer.

To quantitatively assess the conformational differences between the 16S rRNA in the structure obtained from grids having or lacking the extra carbon layer, we produced a molecular model from the cryo-EM map obtained from grids containing the extra carbon layer and we calculated a temperature map (Fig. 3C) with respect to the 30S-Inactivated-high-Mg^2+^ class A particles that were imaged without extra layer of carbon on the grids. We found that the conformation of the 16S rRNA in both structures was very similar, including the alternative folding adopted by the upper domain of helix 44 and 3’ end of the rRNA molecule.

These results revealed that the air-water interface caused uS2 to fall-off. They also demonstrated that the non-canonical folding observed for helix 44 and 3’ end of the 16S rRNA was induced by the exposure of the ribosomal particles to low magnesium concentrations and it occurs independently of the presence of uS2.

### Transition from the inactive to the active state reverts the decoding center to the standard conformation

Elson and colleagues (Zamir et al., 1969, 1971) described that incubation of inactive 30S subunits at 42 °C in the presence of 10-20 mM Mg^2+^ reverts the structural changes induced by low magnesium concentration to an ‘active’ conformation, in which the 30S subunits regain their ability to bind tRNA.

To structurally describe the conformational changes that this incubation triggers, purified 30S-Inactivated-high-Mg^2+^ ribosomal particles were incubated at 42 °C and imaged by cryo-EM. These particles were called 30S-Activated-high-Mg^2+^. Image classification revealed that the activation treatment had transformed all the 30S subunits to adopt a single conformation (Fig. 4A) that closely resembled the structure of the 30S subunit as described by X-ray crystallography (Fig. 1) (Wimberly et al., 2000). The structure refined to a resolution of 3.6 Å (Supplementary Fig. S1) and allowed us to derive a molecular model from the cryo-EM map (Fig. 4B). We then used this model to calculate a temperature map that compared this structure with the crystallographic structure. We observed that helix 44 and rRNA forming the decoding center closely overlapped indicating that the 42 °C incubation treatment reverted the conformation of the functional domain induced by low magnesium concentration to that observed in the 70S ribosome and the 30S subunit structure produced by X-ray crystallography (Fig. 4C).

**Figure 4.**
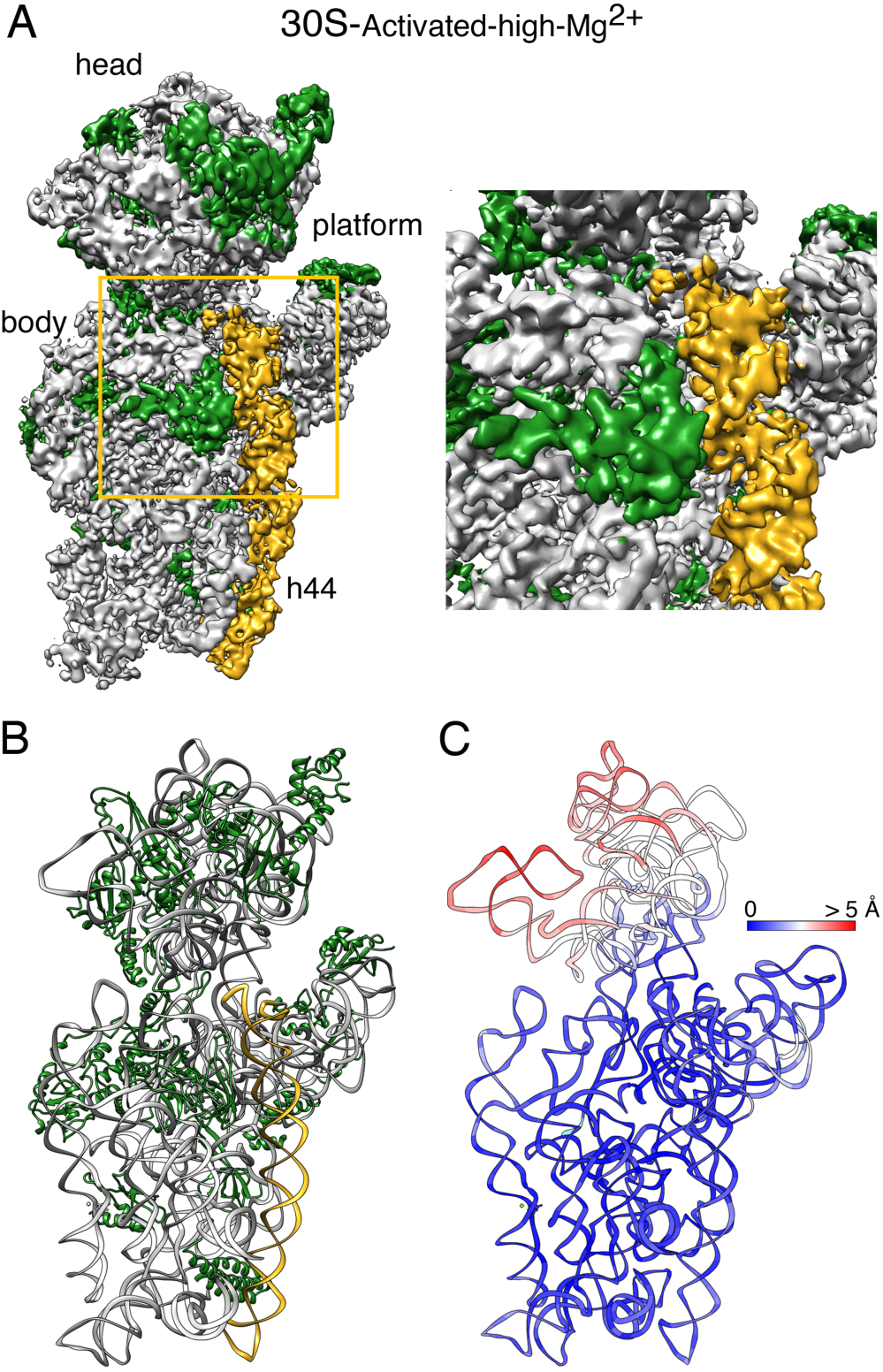
Cryo-EM structure of the 30S-Activated-high-Mg2+ particle. (A) Front view (left panel) of the cryo-EM map obtained for the 30S-Activated-high-Mg2+ particle. The framed area is shown as a zoomed in view in the right panel. Main landmarks of the 30S subunit are labeled. The rRNA is shown in light grey and the r-proteins are shown in green. Helix 44 is shown in golden rod orange. (B) Molecular model of the 30S-Activated-high-Mg2+ particle. The rRNA and the r-proteins are colored as in panel (A). (C) Temperature map of the 30S-Activated-high-Mg2+ molecular model. The rRNA is colored according to the r.m.s.d. deviation (Å) with respect to the structure of the 30S subunit obtained by X-ray crystallography (PDB ID 4V4Q).

### Extended exposure to low magnesium concentration induces unfolding of rRNA structural domains

Next, we inquired about the effect of extended exposure of the 30S subunits to low concentration of magnesium ions. To this end, the purified 30S subunits were maintained in buffer containing 1.1 mM magnesium acetate before imaging them by cryo-EM. Similar classification approaches to those followed in the previous samples revealed that the 30S ribosomal subunits coexisted under low magnesium concentrations in two distinct conformations (Fig. 5A). One of the classes represented 64% of the population and these particles generated a cryo-EM map that refined to 3.4 Å resolution (30S-Inactivated-low-Mg^2+^ class A) (Supplementary Fig. S1). The remaining particles generated a different cryo-EM map that refined to 3.9 Å resolution (30S-Inactivated-low-Mg^2+^ class B) (Supplementary Fig. S1). Both classes significantly diverged from the structure considered as a mature 30S subunit and contained structural features only previously observed in structures of immature 30S subunits (Guo et al., 2013; Jomaa et al., 2011; Leong et al., 2013; Razi et al., 2019). Density representing the entirety of helix 44 was missing in the two cryo-EM maps (Fig. 5A). Similarly, helix 23 and 24 in the platform region exhibited highly fragmented densities and the r-proteins bound to this region (bS6, uS11, bS21 and bS18) were also not observed in the cryo-EM map (Fig. 5B & 5C). The EM grids used to image this sample did not contain an additional layer of continuous carbon and consequently uS2 was missing from the cryo-EM maps of both classes (Fig. 5A, lower panel). The main difference observed between class A and B was in the head domain that was completely missing in the cryo-EM map for class B. Density for all regions of this domain was present in the map obtained for class A, except for r-protein uS7 that similar to other structures or the 30S subunit obtained in solution, may be adopting a flexible conformation (Fig. 5A, top panel).

**Figure 5.**
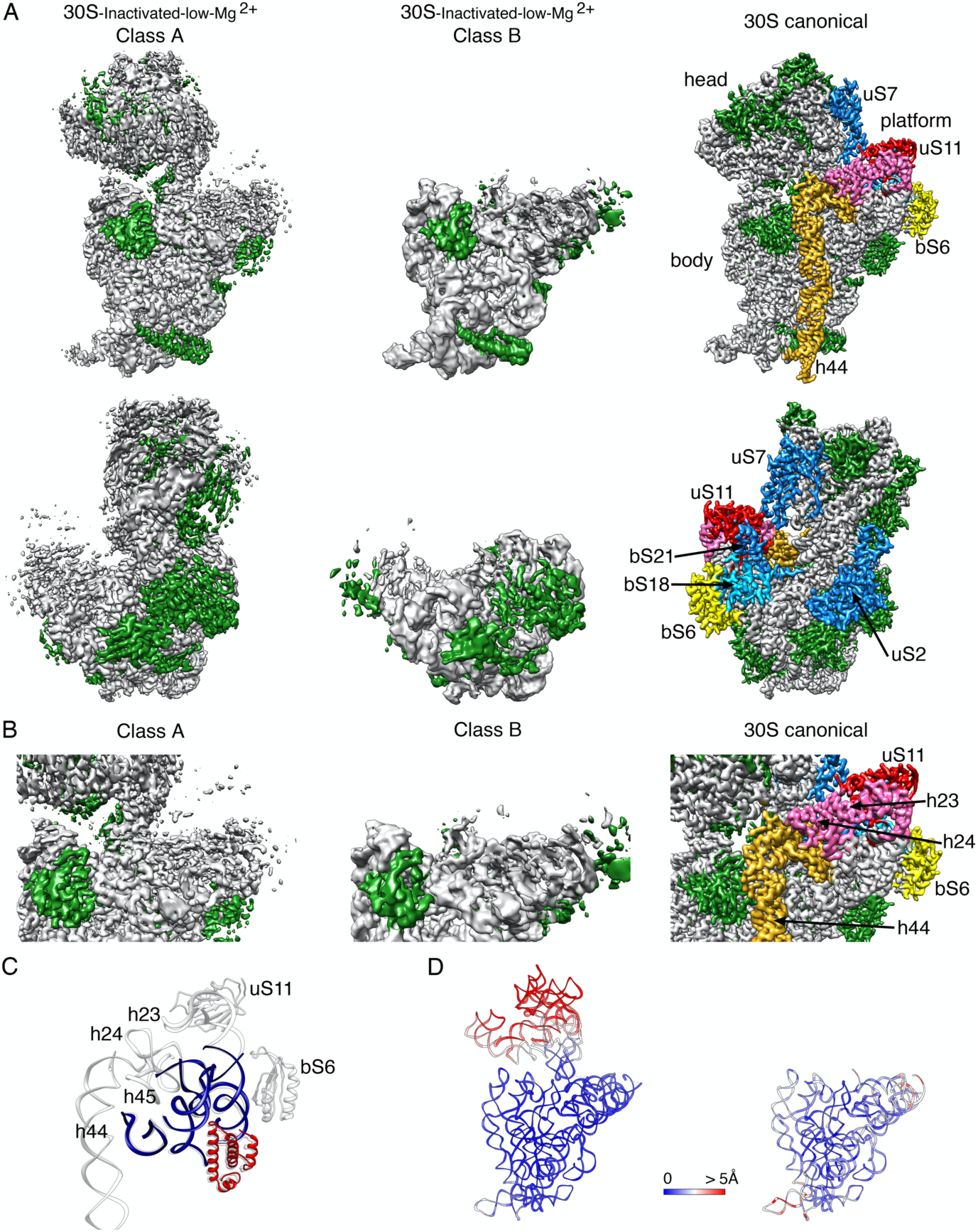
Cryo-EM structure of 30S-Inactivated-low-Mg^2+^ particle. (A) Front (top panels) and back (bottom panels) views of the two conformers, class A and B of the 30SInactivated-low-Mg^2+^ particle. The rRNA is shown in light grey and the r-proteins are shown in green. Helix 44 is shown in golden rod orange. These structures are shown side-by-side with the structure of the 30S subunit obtained by X-ray crystallography (PDB file 4V4Q). In this structure, the rRNA is displayed in light grey, the r-proteins in green and helix 44 in golden rod orange. The r-proteins uS2, bS6, uS7, uS11, uS18 and bS21 for which a representative density does not appear in the cryo-EM maps of the 30S-Inactivated-low-Mg^2+^ particle are indicated using a color different than green. These proteins and other landmarks of the 30S subunit are labeled. (B) Zoomed in view of the decoding and platform region of the cryo-EM maps obtained for the 30S-Inactivated-low-Mg^2+^ class A and B using the same color coding as in (A). (C) Overlap of helix 44 and platform region of the molecular model derived from the cryo-EM structure of the 30S-Inactivated-low-Mg^2+^ class A and the corresponding region from the structure of the 30S subunit obtained by X-ray crystallography. Parts of the structure present in the cryo-EM structure of the 30S-Inactivated-low-Mg^2+^ class A are displayed in navy blue and red. The atomic model of the X-ray structure with all the elements of the complete 30S subunit structure is displayed in light grey. (D) Temperature maps of the 30S-Inactivated-low-Mg^2+^ class A and B molecular models. The rRNA is colored according to the r.m.se.d. deviation (Å) with respect to the structure of the 30S subunit obtained by X-ray crystallography (PDB ID 4V4Q).

The molecular model derived from the 30S-Inactivated-low-Mg^2+^ class A and B cryo-EM maps were used to calculate a temperature map (Fig. 5D) to compare the conformation of the 16S rRNA in these structures with that of the crystallographic structure (Wimberly et al., 2000). The body and platform regions present in the cryo-EM map of the 30S-Inactivated-low-Mg^2+^ class A showed a close overlap. However, we found that the head domain was tilted backwards in class A, similarly to what we observed in the structures derived from the 30S-Inactivated-high-Mg^2+^ particles.

These structures suggest that continued exposure of the mature 30S subunit to low magnesium concentrations destabilizes large structural motifs of the ribosomal subunit, including the central (platform), 3’ major (head) and 3’ minor (helix 44) domains.

### The helix 44 conformation dictates the motions exhibited by the free 30S subunits

We noticed that in all the cryo-EM maps obtained for the 30S subunit, density was clear and mostly complete for the body and platform domain but was fuzzier and fragmented in the head region (Fig. 6A). We interpreted the partial fragmentation of these densities as an indication of the motions that the head region experiences when the 30S subunits are in solution. Using multi-body refinement (Nakane et al., 2018), we investigated these motions of the 30S subunit and whether these motions differ for each one of the conformations of the 30S subunit described in this study. To this end, we split the cryo-EM maps into three bodies corresponding to three of the major domains of the 30S subunit: body, platform and head. After multi-body refinement, we compared the resulting maps obtained for the head domains with those from the consensus refinement. Visual inspection of their central section (Fig. 6) showed a small improvement, particularly in the region furthest away from the center of the map.

**Figure 6.**
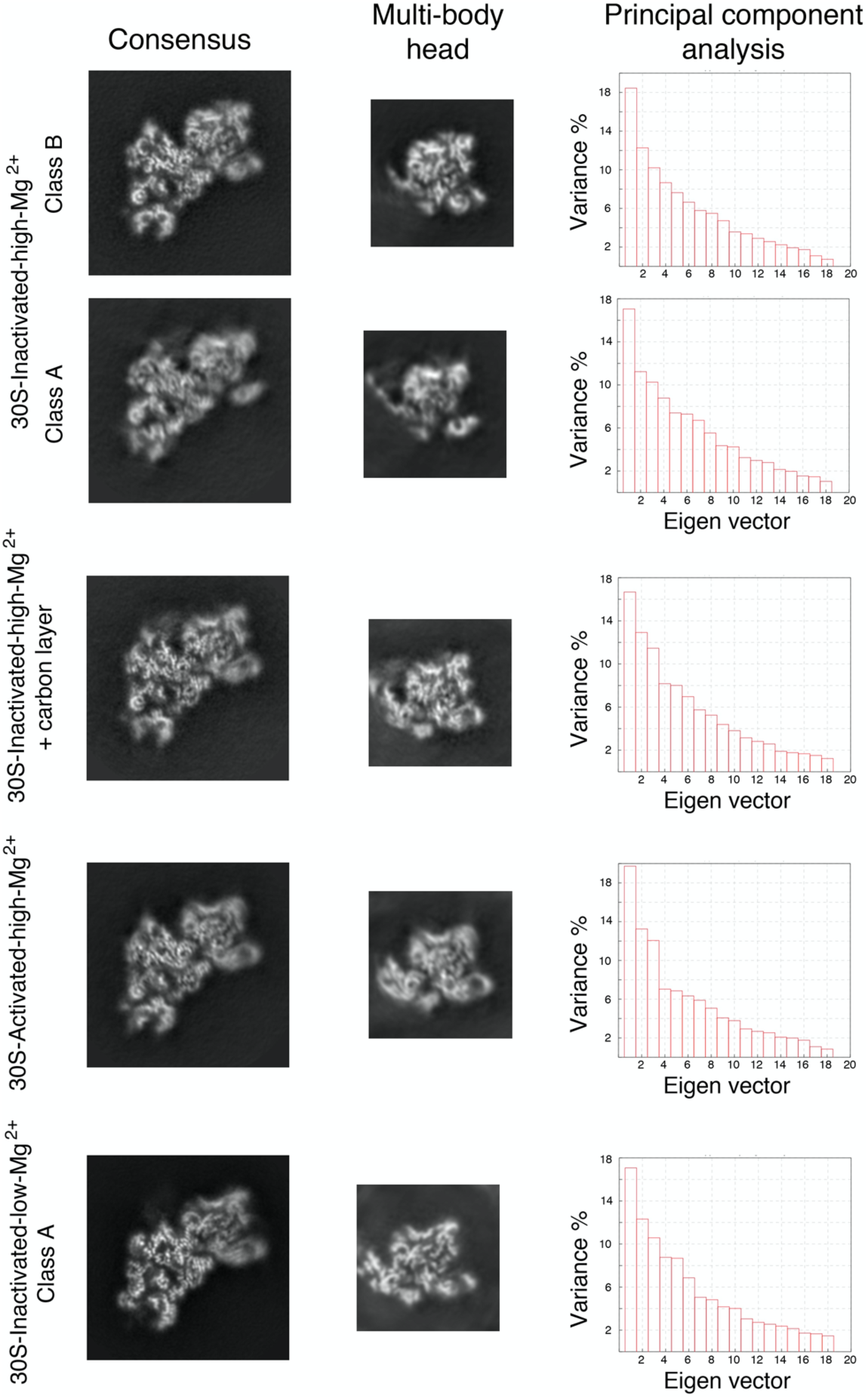
Multibody refinement analysis of the cryo-EM structures. The dataset for each type of 30S particle was analyzed using multibody refinement to visualize their motions. The left and middle panels show respectively the central sections of the cryo-EM maps of the obtained 30S structures after the consensus refinement and of the head after multibody refinement. The high-resolution features of the density maps are apparent in the body region but are slightly blurred in the head domain in the central sections of the maps obtained through consensus refinement. High-resolution features of the head become more apparent in the maps of this region obtained through multibody refinement. The right panels show the principal component analysis of the different structures using the multibody refinement routine in Relion 3.0. This analysis indicated that between 38-43% of the variance is due to the movements of the head with respect to the body and platform.

More importantly, this approach performs a principal component analysis of the variance in the rotations and translations of the three bodies and allowed us to generate movies describing the most important motions in the different 30S subunit populations. In all cases, this analysis revealed that between 38-43% of the variance in the rotations and translations of the three bodies is explained by the first three eigenvectors (Fig. 6). Movies of the reconstructed body densities repositioned along these three eigenvectors revealed that they correspond to motions of the head with respect to the body and platform. Interestingly, the most prevalent of these movements revealed by the first eigenvector was not the same for Jahagirdar *et al*. all 30S populations. We found that the main motion that 30S subunit populations with a helix 44 in the unlatched position undergo is a rotation of the head around an axis longitudinal to the longest dimension of the particle. The other two motions along the second and third eigenvectors are back and forward tilting movements of the head with respect to the body. This is the case for the 30S-Inactivated-high-Mg^2+^ class A (Supplementary Movie 2) and 30SInactivated-Carbon-high-Mg^2+^ (Supplementary Movie 4). In the case of the 30SActivated-high-Mg^2+^ particles, where helix 44 is in the conformation described by the crystal structure, the most frequent motions described by the first two eigenvectors are the tilting movements (Supplementary Movie 5) and the head rotation represents the third most frequent motion. The 30S subunit populations in which part of helix 44 present fragmented densities (30S-Inactivated-high-Mg^2+^ class B) showed only head tilted motions along the first three eigenvectors (Supplementary Movie 3) and head rotation motions were not observed. Particles that belonged to the 30S-Inactivated-low-Mg^2+^ class A and showed complete absence of density for helix 44 exhibited a head tilting motion (Supplementary Movie 6) as the most predominant motion, but the motion along the second eigenvector described a head rotation motion similar to that shown by the 30S subunit populations with a helix 44 in the unlatched position.

Overall, these results indicate that the conformation adopted by helix 44 seems to dictate the most predominant motions exhibited by the free 30S subunits.

## DISCUSSION

The transition of the decoding center in the 30S subunit from the active to the inactive state was one of the first conformational rearrangements discovered in ribosomes (Moazed et al., 1986; Zamir et al., 1969, 1971). Chemical probing experiments at the time revealed that adoption of the inactive conformation involved large-scale changes structural changes in the neck region and the fraction of helix 44 involved in the formation of the decoding center. More recently, RNA SHAPE experiments (McGinnis et al., 2015; McGinnis and Weeks, 2014) proposed an alternative base pairing in the 16S rRNA forming the decoding center in the inactive conformation. In particular, nucleotides 1402-1408, which form an irregular helix at the top of helix 44 pairing with nucleotides 1492-1500 in the conventional structure, undergo a register shift and pair with positions 921-927 within helix 28 and nucleotides 1390-1401 form an unpaired loop.

In this work, we directly visualized using cryo-EM the inactive conformation of the 30S subunit. The obtained high-resolution cryo-EM map showed a structure that diverged from the one proposed in the RNA SHAPE experiments (McGinnis et al., 2015). In the cryo-EM map obtained for the ‘inactive’ conformation nucleotides 1532-1534 at the 3’ end of the 16S rRNA base paired with nucleotides 921-923 in helix 28 causing the unfolding of the bottom of this helix and the top of helix 44. The newly formed base pairs force the 3’ end of the 16S rRNA in a conformation that resembles the conformation adopted by mRNA during translation.

The functional importance of the “inactive” conformation of the 30S subunit has been highlighted in previous studies (Karbstein, 2013; Myasnikov et al., 2009). Multiple translation initiation factors and RNAses affecting 30S subunit turnover bind at sites overlapping with helices 28 and 44, suggesting that the “active-inactive” transition observed by cryo-EM could govern accessibility of these factors and regulate translation initiation and ribosome quality control.

Our study also characterizes the motions that the different 30S subunit subpopulations exhibit when they are in solution and not constrained in a crystal lattice. X-ray crystallographic and cryo-EM structures from many laboratories have described the large-scale conformational changes the ribosome undergoes during protein synthesis (Noeske and Cate, 2012). For example, the movement of the tRNAs from the classical A/A and P/P configurations to the hybrid A/P and P/E states is driven by a rigid body rotation of the small subunit with respect to the large subunit by 10° around an axis perpendicular to the subunit interface (ratchet-like motion) (Frank and Agrawal, 2000). This movement is also accompanied with a simultaneous swiveling of the 30S subunit head domain in the direction of the E site (Spahn et al., 2004). Swiveling motions of the head also accompany the process of termination and ribosome recycling (Dunkle et al., 2011; Yokoyama et al., 2012). The head domain also exhibits other motions linked to other physiological processes such as hibernation. Binding of ribosome modulation factor or hibernation promoting factor triggers a displacement of the head away from the central protuberance (backward tilting), promoting formation of 100S ribosome dimers (Polikanov et al., 2012). All these motions have been observed in the context of the 70S ribosome. In our study, the use of cryo-EM combined with multibody refinement processing approaches (Nakane et al., 2018) allowed us to investigate the most prominent motions of the different conformational subpopulations identified for the 30S subunit. Overall, we found that the motions of swiveling and backward tilting that the head domain of the 30S subunit undergoes in the context of the 70S ribosome are also exhibited by the free 30S subunits. Our results suggest that the conformation adopted by helix 44 in a particular 30S subunit populations is the main determinant dictating the most predominant motion.

An important motivation for this study was to provide a more extensive comparative reference for those studies on assembly of the 30S subunit that use single deletion or depletion strains of particular assembly factors. Drawing meaningful conclusions from the heterogeneous mixture of ribosomal particles that these strains accumulate is only possible when a comprehensive description of the conformations adopted by the mature subunit is available. Only then it is possible to separate from the observed pool of structures, those truly representing immature assembly states and derive information about the function of specific assembly factors.

Current sample preparation techniques in cryo-EM suffers from a fundamental problem. Macromolecules in solution drift during vitrification and get exposed to the air-water interface at the top and bottom of the thin films formed upon blotting. This exposure is damaging to the macromolecular assemblies and can pull their components apart and destroy them. Our results show that even for specimens considered resilient such as the ribosome or the 30S subunit, current cryo-EM sample preparation methods still cause damages. These results will contribute to differentiate bona fide structural defects observed on heterogeneous mixtures of ribosomal particles from those caused by the sample preparation technique.

Overall, this study provides new insights into the alternative conformations and motions adopted by the 30S ribosomal subunit. The existence of one of them, named by Elson as the “inactive” conformation, was discovered over five decades ago. It has been only now with the recent improvements in electron microscopes and direct electron detector cameras that we have been able to describe these structures at high-resolution.

## MATERIAL AND METHODS

### Cell strains

We used the *Escherichia coli* K-12 (BW25113) strain from the Keio collection (Baba et al., 2006) to produce the purified 30S subunits used in this study.

### Purification of 30S ribosomal particles

To obtain purified 30S subunits, we first produced a 30 mL saturated overnight culture of *E. coli* K-12 strain (BW25113) in LB media and this culture was used to inoculate a 3 L culture of LB media. Cells were grown at 37 °C with shaking at 225 rpm in an Innova 44 incubator shaker (New Brunswick) until they reached an OD_600_ of 0.6. Harvesting of the cells was done by centrifugation at 3,700 *g* for 15 min and the obtained pellets were chilled to 4 °C and all the subsequent steps were done at this temperature. These pellets were resuspended in 7 mL of buffer A (20 mM Tris-HCl at pH 7.5, 10 mM magnesium acetate, 100 mM NH_4_Cl, 0.5 mM EDTA, 3 mM 2-mercaptoethanol, and a protease inhibitor mixture (cOmplete Protease Inhibitor Mixture Tablets; Roche) and DNaseI (Roche). Cell lysis was done by passing the cell suspension through a French pressure cell at 1,400 kg/cm^2^ three consecutive times and the cell debris were cleared by spinning the lysate at 59,000*g* for 30 min. The supernatant was layered over a sucrose cushion of equal volume composed of 37.6% sucrose in buffer B (20 mM Tris-HCl pH 7.5, 10 mM magnesium acetate, 500 mM NH_4_Cl, 0.5 mM EDTA, and 3 mM 2-mecaptoethanol), and then spun down for 4.5 hours at 321,000 *g*. The pellet containing the ribosomal particles was resuspended in buffer C containing 10 mM Tris-HCl pH 7.5, 10 mM magnesium acetate, 500 mM NH_4_Cl, 0.5 mM EDTA and 3 mM 2-mecaptoethanol and then spun for 16 hours at 100,000 *g*. This produced a pellet containing washed ribosomal particles that was resuspended in buffer F (10 mM Tris-HCl, pH 7.5, 1.1 mM magnesium acetate, 60 mM NH_4_Cl, 0.5 mM EDTA, and 2 mM 2-mercaptoethanol). To separate the fraction containing the 30S subunits, approximately 120 A_260_ units of resuspended crude ribosomes were then applied to 34 mL of 10–30% (wt/vol) sucrose gradients prepared with buffer F. The gradient was centrifuged for 16 hours at 40,000 *g* on a Beckman Coulter SW32 Ti rotor and fractionated using a Brandel fractionator apparatus and an AKTA Prime FPLC system (GE Healthcare). Fractions containing the 30S subunits were selected and pooled together based on the UV absorbance at A_254_. Subsequently, they were spin down for another 4.5 hours at 321,000*g* on a Beckman 70Ti rotor. The pellet was resuspended differently according to the final conditions we intended to study the 30S subunit. The 30S subunits under ‘low Mg^2+^ conditions’ (30SInactivated-low-Mg^2+^) were resuspended back in buffer F, whereas the 30S subunits under ‘high Mg^2+^ conditions’ (30SInactivated-high-Mg^2+^ and 30SInactivated-Carbon-high-Mg^2+^) and were resuspended in buffer E (10 mM Tris-HCl, pH 7.5, 10 mM magnesium acetate, 60 mM NH_4_Cl and 3 mM 2-mecaptoethanol). ‘Activated’ 30S subunits (30SActivated-high-Mg^2+^) were resuspended in modified buffer E with a different concentration of magnesium acetate (20 mM) and stored at −80 °C. To perform the activation, these 30S subunits were heated at 42 °C for 20 minutes and then diluted by mixing equal volumes of the sample with buffer E not containing any magnesium acetate.

### Cryo-electron microscopy

Sample vitrification was performed using a Vitrobot (Thermo Fisher Scientific Inc.). For all samples a volume of 3.6 μL of the diluted sample was applied to holey carbon grids (c-flat CF-2/2-2C-T) previously washed in chloroform for two hours and treated with glow discharged in air at 5 mA for 15 seconds right before the sample was applied. For the 30SInactivated-Carbon-high-Mg^2+^ sample, we used holey carbon grids containing an additional layer of continuous thin carbon (5-10nm). The concentration of ribosomal particles in the solution applied to the grid varied between 200-300 nM depending on the sample. In the Vitrobot, each grid was blotted once for 3 seconds and with a blot force +1 before they were plunged into liquid ethane. The Vitrobot was set at 25 °C and 100% relative humidity.

Data acquisition was performed using EPU software at FEMR-McGill using a Titan Krios microscope at 300 kV equipped with a Falcon II direct electron detector (Thermo Fisher Scientific Inc.). Movies for the 30S-Inactivated-high-Mg^2+^, 30S-Inactivated-Carbon-high-Mg^2+^, 30S-Activated-high-Mg^2+^ and 30S-Inactivated-low-Mg^2+^ datasets were collected with a total dose of 50, 52, 52 and 50 e^-^/Å^2^, respectively. All datasets except for the 30S-Inactivated-Carbon-high-Mg^2+^ particles were collected as movies with seven frames acquired in 1 second exposure at a magnification of 75,000x, producing images with a calibrated pixel size of 1.073 Å. Movies for the 30S-Inactivated-Carbon-high-Mg^2+^ particle were collected in the same manner but with 30 frames. The nominal defocus range used during data collection was between −1.25 to −2.75 μm.

### Image processing

Collected movies were corrected for beam-induced motion using Relion’s implementation of the MotionCor 2 algorithm (Zheng et al., 2017; Zivanov et al., 2018). Correction for the Contrast Transfer Function (CTF) was done using the Gctf program (Zhang, 2016). Subsequent processing steps were done using Relion-3 (Zivanov et al., 2018). Particle images were selected and extracted from the micrographs using auto-picking and subsequently subjected to one or more cycles of reference-free 2D classification to remove false positive and damaged particles. This process produced clean datasets for the 30S-Inactivated-high-Mg^2+^, 30SInactivated-Carbon-high-Mg^2+^, 30SActivated-high-Mg^2+^ and 30SInactivated-low-Mg^2+^ samples comprised of 565,255, 334,903, 407,623 and 658,065 particles, respectively. These datasets were used for 3D classifications to separate the different conformational subpopulations existing in each sample. The initial 3D reference used for these classifications was either a 60 Å low pass filtered map of the mature 30S subunit created from 4V4Q.pdb (Schuwirth et al., 2005) using the Xmipp program (de la Rosa-Trevin et al., 2013) or the intermediate cryo-EM maps obtained during classification. No mask was used for the 3D classifications. All 2D classification and 3D classification jobs were performed using particle images binned by 4 and a pixel size of 4.292 Å. In each dataset, maps obtained in the classification steps were visually inspected in Chimera (Pettersen et al., 2004) and particles assigned to maps representing the same conformation were pooled together and used for refinement and produce high resolution maps. A soft-mask was applied to all refinements. These masks were created with ‘relion_mask_create’ command extending the binary mask by four pixels and creating a soft-edge with a width of four pixels. The initial threshold for binarization of the mask varied depending on the structure. As initial map for the refinement procedures we used either a 60 Å low pass filtered map of the mature 30S subunit created from 4V4Q.pdb (Schuwirth et al., 2005) using the Xmipp program (de la Rosa-Trevin et al., 2013) or the cryo-EM maps obtained during classification after they were re-scaled back to full-size with a pixel size of 1.073 Å. Sharpening of the final cryo-EM maps and the local analysis was done with Relion (Zivanov et al., 2018).

### Isotropic map normalization

The cryo-EM maps for 30S-Inactivated-high-Mg^2+^ class A and 30S-Activated-high-Mg^2+^ were slightly affected by directional resolution anisotropy caused by the presence of preferential specimen orientations. In these cases, the underrepresented macromolecular views result on low map amplitudes along these directions causing a stretching in the reconstructed volume. To the best of our knowledge, there is not currently any computational tool in single particle analysis to reduce the effect of preferential specimen orientations in the reconstructed map other than tilting the specimen (Tan et al., 2017). In this work, we compensate the map stretching caused by preferential specimen orientations by an isotropic amplitude map normalization in Fourier space. In this approach, we normalize map amplitudes at a certain resolution along all possible map directions for resolutions higher than 9-10 Å. At these resolution ranges, respective Fourier components encode the information of secondary structures presented in the map. Our hypothesis is that these secondary structures should be oriented approximately random wise. Then, the amplitudes of map Fourier components should be approximately isotropic for resolutions higher than 9-10 Å. Our map restoration method follows these steps: (1) The Fourier transform of the cryo-EM map is obtained; (2) Starting from 9-10 Å, a shell of the map Fourier transform is extracted; (3) The map amplitudes within the extracted shell and the respective q=75% percentile value is obtained; (4) Map amplitudes at the extracted shell are modify so their value is set to q; (5) A new shell at higher resolution is extracted and steps (1) to (4) are repeated; (6) This process is iterated until the shell at highest resolution (Nyquist resolution) is processed; (7) The inverse Fourier transform of the isotropic amplitude normalized map is computed obtaining a new corrected map. This map restoration approach transforms map Fourier amplitudes only, without modifying map phases.

### Multibody refinement and motion analysis

We used the Relion-3 implementation (Nakane et al., 2018) to perform the multibody refinement and motion analysis. The same approach was followed for all 30S subunit populations. The consensus cryo-EM map obtained for each class was divided into three bodies corresponding to the three major domains of the subunit: body, platform and head (Fig. 6B). The masks for the corresponding bodies were made using a 25 Å low-pass filtered version of the consensus map. We use available atomic models of the 30S subunit (PDB ID 4V4Q) to determine the boundaries between the bodies. By extending the binary map by 10 pixels and placing a soft edge with of 10 pixels, all three bodies overlapped with each other. The multibody refinement job was run with downsized particle images with a box size of 218 and a pixel size of 1.496 Å. The standard deviation of the Gaussian prior on the rotations was set to 10 degrees for all three bodies, and the standard deviations on the body translations were all set to 2 pixels. Based on the domain architecture of the 30S subunit, the head was set to rotate with respect to platform and body. Multibody refinement was started using an initial angular sampling rate of 1.8 degrees, an initial offset range of 3 pixels and an initial offset step of 0.25 pixels.

### Map analysis and Atomic model building

Before start building the molecular models for each structure, the obtained cryo-EM maps from Relion, the connectivity of the densities of the cryo-EM maps was improved using automatic sharpening as implemented in the PHENIX suite (Adams et al., 2010). Model building of all maps started by fitting the 4V4Q.pdb (Schuwirth et al., 2005) X-ray structure of the 30S subunit into the obtained cryo-EM maps using the rigid-body docking tools in Chimera (Pettersen et al., 2004). The molecular models were then built through multiple rounds of manual model building in Coot (Emsley and Cowtan, 2004; Emsley et al., 2010) and real space refinement using Phenix (Adams et al., 2010).

Panel for figures were prepared using Pymol program (The PyMOL Molecular Graphics System, Version 2.3.3, Schrodinger,LLC), UCSF Chimera and Chimera X. Figures were assembled using Photoshop (Adobe). Secondary structures of RNA were produced with DSSR (http://x3dna.org) (Lu and Olson, 2008) and Varna (http://varna.lri.fr) (Darty et al., 2009).

## DEPOSITED MAPS AND ASSOCIATED COORDINATES

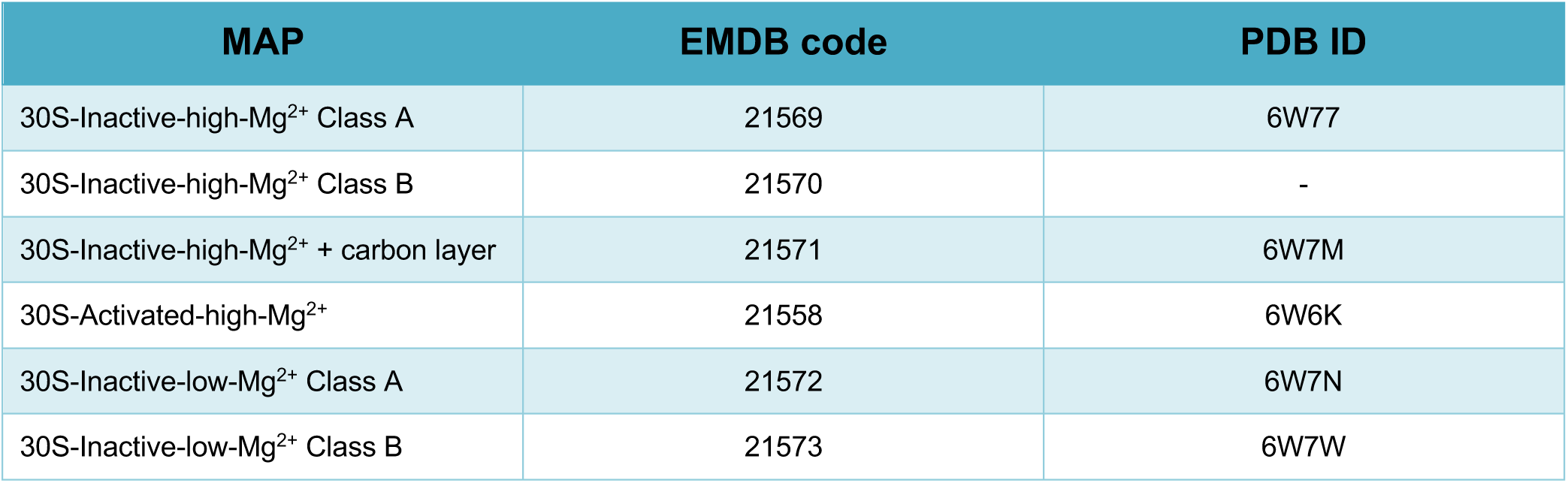

### ADDITIONAL FILES

Supplementary File 1. Supplementary Figure S1: Resolution analysis of the cryo-EM maps. Supplementary Movies Captions.

Supplementary Table S1. Cryo-EM data acquisition, processing and map and model statistics.

## ACKNOWLEDGEMENTS

We thank Kelly Sears, Mike Strauss and other staff members of the Facility for Electron Microscopy Research (FEMR) at McGill University for help with microscope operation and data collection. We acknowledge Clara Ortega for assistance with graphic design.

