## Supplementary Fig. for "Alternative Conformations and Motions Adopted by 30S Ribosomal Subunits Visualized by Cryo-Electron Microscopy"

**This supplement contains:**

Supplementary Figure S1

Supplementary Movies Captions

Supplementary Table S1

### SUPPLEMENTARY FIGURE

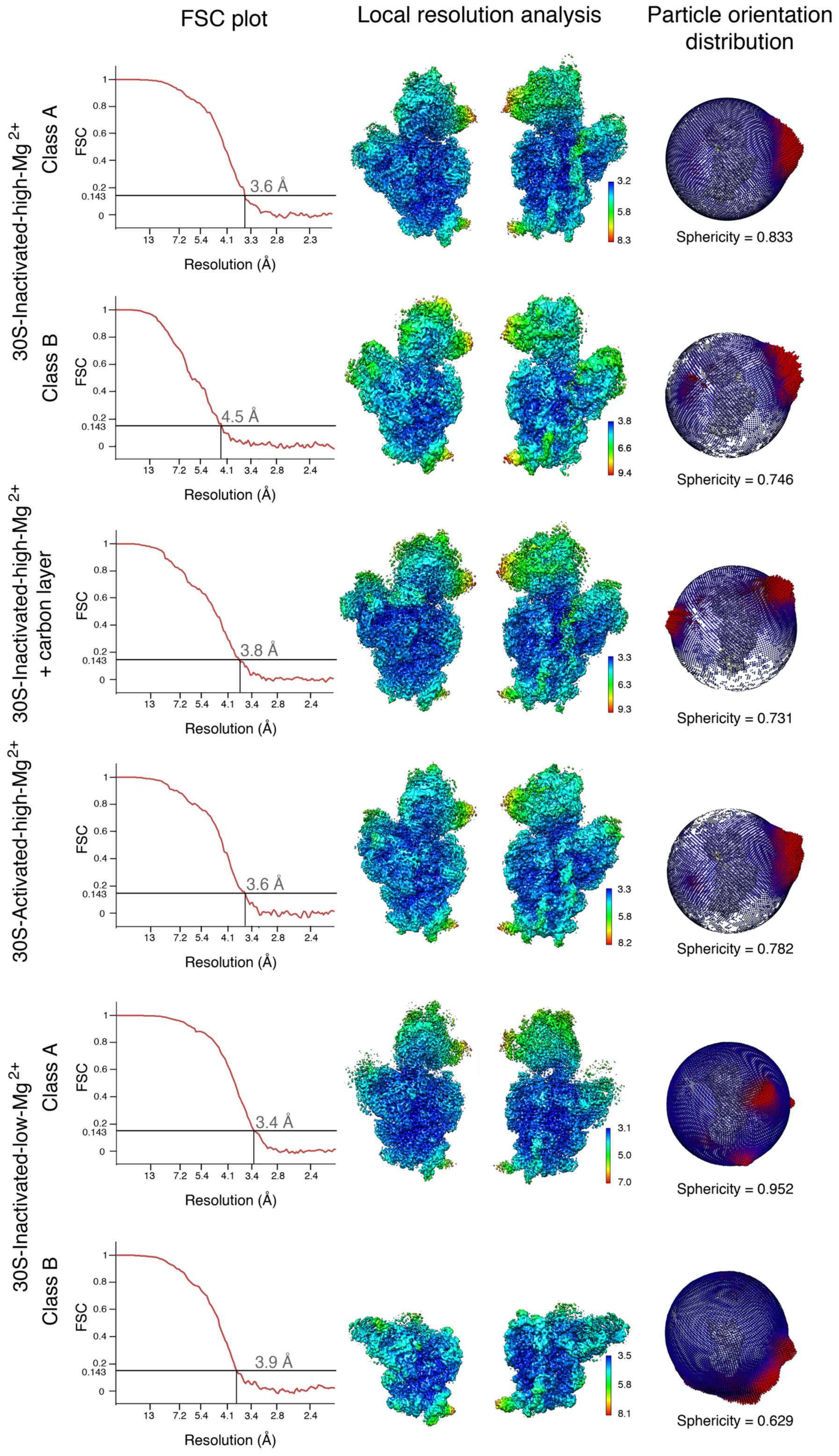

**Supplementary Figure S1.** Resolution analysis of the cryo-EM maps. Left column shows the Fourier shell correlation (FSC) plots for the different cryo-EM maps obtained in this study. Resolution is reported using FSC threshold of 0.143. The middle column displays the cryo-EM maps colored according to the local resolution analysis performed with Relion. The right column shows a sphere representing the angular distribution of the particles in each dataset. Each view angle is represented as a dot in the sphere. The height and color of the sphere relate to the number of particles representing each view. Areas in red indicate a higher number of particles representing those particular views.

### SUPPLEMENTARY MOVIE CAPTIONS

**Supplementary Movie 1.** Conformational transition between the canonical structure of the 30S subunit obtained by X-ray crystallography and the 30S-$_{\text{Inactivated-high-Mg}^{2+}}$ particle. The transition between both conformations involves repositioning of nucleotides 1532-1534 that are not forming any base pairing in the conventional structure to base pair with nucleotides 921-925. In this process the bottom of helix 28 (region formed by nucleotides 1391-1396 and 921-925) and top of helix 44 formed in the canonical structure by nucleotides 1397-1407 and 1494-1503 become unfolded. The 3' end of the 16S rRNA (distal to nucleotide 1534) adopts a conformation similar to that adopted by mRNA during translation. The rRNA is display in light grey, the r-proteins in red. Helices 28, 44 and 45 are colored in navy blue except the region on these helices that unfolds during the conformational change that is colored in yellow. Nucleotides 1532-1534 that upon the conformational transition form base pares with nucleotides 921-925 are shown in red.

**Supplementary Movie 2-6.** Main motions of the 30S particles. Each movie shows the main motion represented by the first eigen vector in each type of 30S particle.

### SUPPLEMENTARY TABLE

**Supplementary Table S1.** Cryo-EM data acquisition, processing and map and model statistics.

| | | 30S-Inactive-high-$Mg^{2+}$ Class A | 30S-Inactive-high-$Mg^{2+}$ Class B | 30S-Inactive-high-$Mg^{2+}$ +carbon layer | 30S-Activated-high-$Mg^{2+}$ | 30S-Inactive-low-$Mg^{2+}$ Class A | 30S-Inactive-low-$Mg^{2+}$ Class B |
|---|---|---|---|---|---|---|---|
| **Data collection** | | | | | | | |
| Microscope | | Titan Krios | | Titan Krios | Titan Krios | Titan Krios | |
| Detector | | Falcon II | | Falcon II | Falcon II | Falcon II | |
| Nominal Magnification | | 75,000x | | 75,000x | 75,000x | 75,000x | |
| Voltage (kV) | | 300 | | 300 | 300 | 300 | |
| Total exposure ($e^-/Å^2$) | | 50 | | 52 | 52 | 50 | |
| Defocus range (μm) | | -1.25 to -2.75 | | -1.25 to -2.75 | -1.25 to -2.75 | -1.25 to -2.75 | |
| Calibrated physical pixel size (Å/px) | | 1.073 | | 1.073 | 1.073 | 1.073 | |
| **Reconstruction and refinement** | | | | | | | |
| Particles | | 446,530 | 118,725 | 334,903 | 407,603 | 421,738 | 236,327 |
| Map sharpening B factor | | -116 | -136 | -98 | -100 | -121 | -161 |
| Resolution (Å) | | 3.6 | 4.5 | 3.8 | 3.6 | 3.4 | 3.9 |
| FSC Threshold | | 0.143 | 0.143 | 0.143 | 0.143 | 0.143 | 0.143 |
| **Model composition** | | | | | | | |
| RNA chains | | 1 | - | 1 | 1 | 1 | 1 |
| Protein chains | | 17 | - | 20 | 17 | 14 | 9 |
| **Model Building** | | | | | | | |
| Protein Geometry | Poor rotamers | 0.00% | - | 0.1% | 0.00% | 0.00% | 0.12% |
| | Favored rotamers | 95.47% | - | 93.90% | 96.08% | 95.94% | 96.71% |
| | Ramachandran outliers | 0.21% | - | 0.31% | 0.00% | 0.31% | 0.00% |
| | Ramachandran favored | 93.73% | - | 92.28% | 93.07% | 91.60% | 95.92% |
| | Cβ deviations >0.25Å | 0.00% | - | 0.00% | 0.00% | 0.00% | 0.00% |
| | Bad bonds | 0.00% | - | 0.00% | 0.00% | 0.00% | 0.00% |
| | Bad angles | 0.00% | - | 0.00% | 0.00% | 0.00% | 0.00% |
| Nucleic Acid Geometry | Probably wrong sugar puckers | 0.86% | - | 1.12% | 0.65% | 0.58% | 0.46% |
| | Bad backbone conformations | 19.42% | - | 27.38% | 20.08% | 16.76% | 20.90% |
| | Bad bonds | 0.00% | - | 0.00% | 0.00% | 0.00% | 0.00% |
| | Bad angles | 0.03% | - | 0.01% | 0.00% | 0.01% | 0.00% |
| Low-resolution criteria | CaBLAM outliers | 4.4% | - | 4.4% | 5.4% | 6.0% | 3.2% |
| | CA Geometry outliers | 0.96% | - | 0.94% | 1.28% | 1.00% | 0.99% |
| Additional validations | Chiral Volumes outliers | 0/9933 | - | 0/10342 | 0/9855 | 0/8827 | 0/5606 |
